# Adjuvant-dependent impacts on vaccine-induced humoral responses and protection in preclinical models of nasal and genital colonization by pathogenic Neisseria

**DOI:** 10.1101/2024.09.07.611809

**Authors:** Epshita A. Islam, Jamie E. Fegan, Joseph J. Zeppa, Sang K. Ahn, Dixon Ng, Elissa G. Currie, Jessica Lam, Trevor F. Moraes, Scott D. Gray-Owen

## Abstract

*Neisseria gonorrhoeae,* which causes the sexually transmitted infection gonorrhea and *N. meningitidis*, a leading cause of bacterial meningitis and septicemia, are closely related human-restricted pathogens that inhabit distinct primary mucosal niches. While successful vaccines against invasive meningococcal disease have been available for decades, the rapid rise in antibiotic resistance has led to an urgent need to develop an effective gonococcal vaccine. Several surface antigens are shared among these two pathogens, making cross-species protection an exciting prospect. However, the type of vaccine-mediated immune response required to achieve protection against respiratory versus genital infection remains ill defined. In this study, we utilize well established mouse models of female lower genital tract colonization by *N. gonorrhoeae* and upper respiratory tract colonization by *N. meningitidis* to examine the performance of transferrin binding protein B (TbpB) vaccines formulated with immunologically distinct vaccine adjuvants. We demonstrate that vaccine-mediated protection is influenced by the choice of adjuvant, with Th1/2-balanced adjuvants performing optimally against *N. gonorrhoeae,* and both Th1/2-balanced and Th2-skewing adjuvants leading to a significant reduction in *N. meningitidis* burden. We further establish a lack of correlation between protection status and the humoral response or bactericidal titre. Combined, this work provides supports the feasibility for a single vaccine formulation to achieve pan-neisserial coverage.

## Introduction

*Neisseria gonorrhoeae* (gonococcus, Ngo) and *Neisseria meningitidis* (meningococcus, Nme) are closely related Gram-negative bacteria that cause two vastly distinct diseases in humans. Ngo is a non-encapsulated sexually transmitted pathogen that primarily inhabits the urogenital mucosa, though also has the capacity to colonize pharyngeal tissues. Clinical outcomes for gonococcal infection can range from no symptoms to an intense inflammatory response in the affected tissues [1]. Without early and appropriate treatment, permanent scarring of the reproductive organs and infertility can occur [2, 3]. Increasing concerns of treatment failure due to the emergence of multidrug resistant Ngo has made the development of a gonococcal vaccine imperative [4, 5]. Ideally, an effective vaccine would provide protection at the mucosal surface to prevent colonization, disease progression and transmission. However, a major impediment in vaccine development efforts is a gap in understanding of the type of immune response needed to achieve protection at these different mucosal sites.

Unlike Ngo, Nme is usually encapsulated and colonizes the upper respiratory tract, typically without symptoms. In rare instances Nme can disseminate into the bloodstream and cause invasive meningococcal disease where meningitis and septicemia can lead to devastating outcomes [6]. The rapidly fatal nature of systemic Nme infections has led to the development of conjugate-capsular vaccines against serogroups A, C, Y and W, and protein-based vaccines to cover serogroup B [7]. Nme vaccines provide remarkable protection against invasive disease and serum bactericidal activity (SBA), a measure of antibody-mediated complement killing of the meningococcus, serves as a well-established correlate of protection [8]. While the capsule-based vaccines have led to a reduction in nasopharyngeal carriage, the serogroup B vaccines do not appear to do so [9–17]. Whether bactericidal antibodies play a key role during mucosal protection also remains undefined.

Considering the high degree of homology between these pathogens, including the prediction that over fifty putative outer membrane proteins may be shared between Ngo and Nme, achieving cross-species protection with antigens derived from either species is highly plausible [18, 19]. Indeed, meningococcal group B outer membrane vesicle (OMV)-based vaccines have been implicated in cross-protection against Ngo and is currently being recommended in the UK for prevention of gonorrhea in high risk populations [20–22]. Conversely, several prospective gonococcal protein antigens are shared with Nme, including receptors involved in nutrient acquisition (TbpAB, LbpAB, TdfF, TdfH, TdfJ, MetQ) and host cell adhesion (NHBA) [23–30]. Hence, examining the efficacy of prospective vaccine antigens against both Ngo and Nme at the preclinical stage is crucial for understanding overall vaccine impact against pathogenic Neisseria.

Fortunately, even though Ngo and Nme are human-specific pathogens with no natural animal reservoirs, mouse models that allow mucosal colonization are available and can serve as useful tools for preclinical assessment. Hormone and antibiotic treated female mice of certain genetic backgrounds support Ngo in the lower genital tract for up to three weeks [31]. Transgenic mice containing human CEACAM1, which is the host surface receptor for Ngo and Nme, enable robust nasopharyngeal colonization of Nme [32]. These models have been vital for the evaluation of existing and experimental vaccines [21, 24, 27, 32–35], as well as shedding light on mechanisms that clear primary infection and protect against reinfection. For Ngo, the development of antibodies and IFNγ-producing Th1 cells during IL-12-enhanced primary infection was shown to be protective against reinfection of the lower genital tract [36]. For Nme, vaccine-mediated antibody-mediated opsonophagocytosis by PMNs has been identified as a key effector mechanism that provides protection against repeat infectious challenge in the nasopharynx [37]. These advances make it enticing to consider whether the same effector mechanisms would be required for protection via parenteral immunization.

Vaccine adjuvants play a crucial role in enhancing and shaping the immune response against the target antigen. Depending on its composition, the adjuvant not only stabilizes and prolongs antigen persistence at the site of injection, but also directly controls the early immune response through activation of specific pathways and recruitment of immune cells [38–40]. While past studies with Ngo and Nme in mice insinuate that a Th1 response would be crucial for mucosal protection, no comparative studies have yet examined the impact of vaccination with different Th-skewing formulations.

In this study we utilize a panel of 5 adjuvants in conjunction with purified soluble recombinant non-lipidated transferrin binding protein B (TbpB), a lipoprotein anchored to the surface of both *Neisseria* species to facilitate iron acquisition from the host protein transferrin. Adjuvants selected for this study include: (1) Alhydrogel, an Alum salt based formulation found in several commercial vaccines, which elicits a predominantly Th2 response[40, 41]; (2) Alhydrogel plus the Th-1 skewing monophosphoryl lipid A (MPL, a detoxified version of *Salmonella enterica* endotoxin, a TLR-4 agonist), a combination that is used in a commercial human papillomavirus vaccine, for a balanced Th1/2 response [42–44]; (3) Addavax, an oil-in-water emulsion of squalene that is the research-grade version of MF59 found in a commercial influenzae vaccine, which elicits a Th2-skewed response [45–47]; (4) Emulsigen D, a veterinary oil-in-water emulsion adjuvant that additionally contains T cell activating nanoparticle dimethyldioctadecyl ammonium bromide (DDA), which is reported to yield a balanced Th1/2 response [48]; and (5) Th1-skewing Complete Freund’s Adjuvant (CFA), which consists of mineral oil and inactivated mycobacteria as the first dose, followed by Th2-biased Incomplete Freund’s Adjuvant (IFA), which contains only mineral oil [49]. When formulated with antigen, CFA and IFA form water-in-oil emulsions.

Harnessing adjuvants characterized by markedly diverse chemical and immunological properties, we hypothesize a nuanced variation in vaccine-driven immune responses. By including both Ngo and Nme colonization models in this study, we methodically examine whether a common set of adjuvant(s) can offer optimal protection against both pathogenic species/sites. Given that humoral responses are used as a standard metric in vaccine development and that monoclonal antibodies against lipooligosaccharide (LOS) have been implicated in gonococcal clearance [35], we further delve into the correlation between humoral signatures and colonization status in matched samples. Overall, the goal of this study is to provide valuable insight into the feasibility of a universally effective protein vaccine and set the stage for unraveling the specific nature of the immune response requisite for attaining optimal mucosal protection.

## RESULTS

### Unique humoral responses induced by TbpB formulated with different adjuvants

To understand what impact adjuvants can have on the immunological response to TbpB vaccines, C57BL/6 mice were immunized with purified recombinant TbpB protein alone or formulated with either Alum, Alum+MPL, Emulsigen D, Addavax or CFA/IFA. Three doses were administered 21 days apart, and serum sampled 14-21 days after each dose to monitor TbpB-specific antibody levels by ELISA. Compared to unvaccinated controls, all the cohorts that received adjuvanted TbpB vaccines produced significantly higher anti-TbpB antibodies after 3 doses (**Figure 1A**), with a dose-dependent anti-TbpB IgG response always apparent. Purified protein alone did not elicit significant levels of IgG, indicating that adjuvants are needed for an efficient humoral response in this model. Total anti-TbpB IgG varied among the different adjuvant groups, with Emulsigen D and Addavax eliciting the highest and lowest titres respectively.

**Figure 1:**
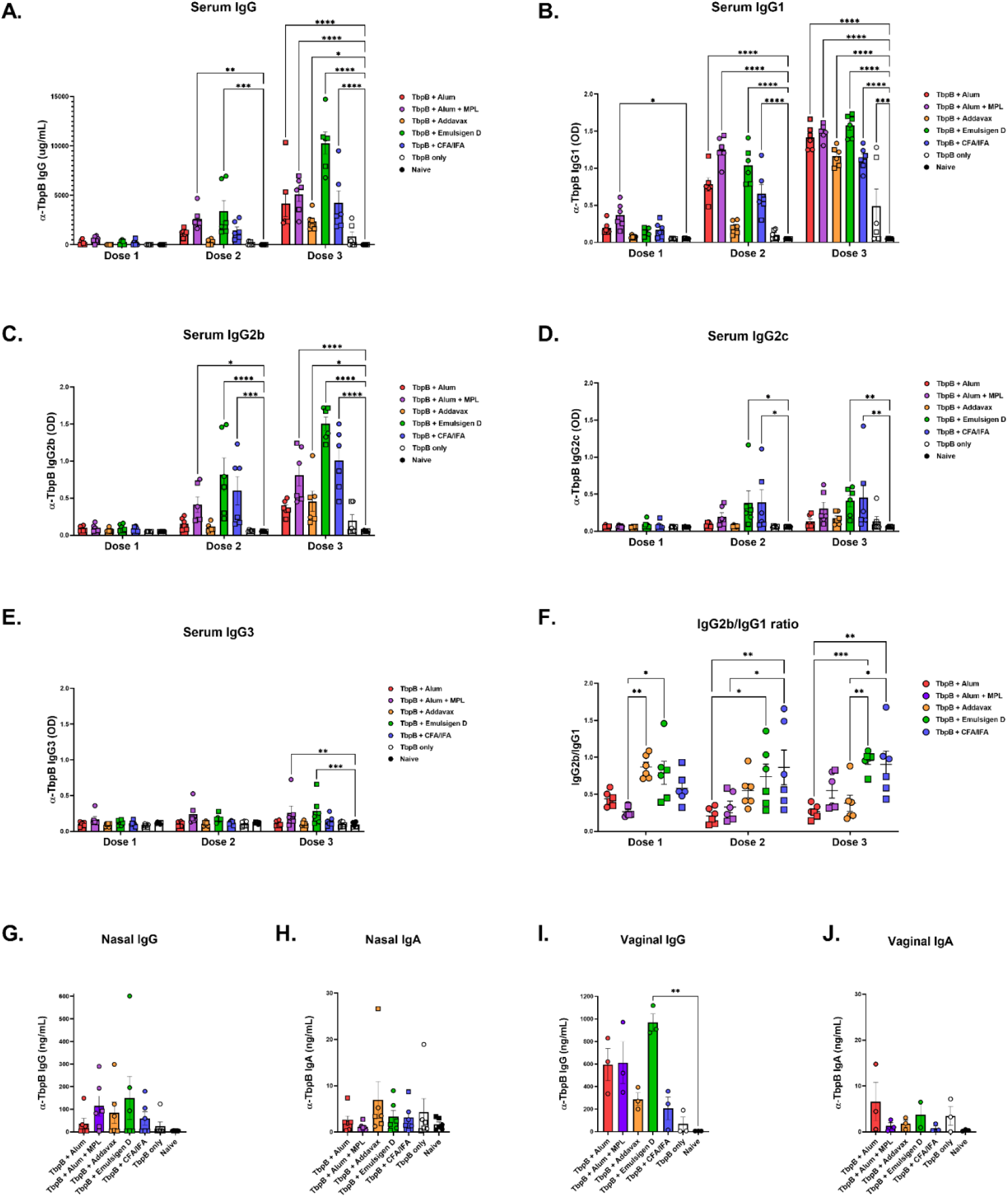
Humoral response to purified TbpB-based vaccines formulated with different adjuvants. **A:** Total anti-TbpB IgG in serum after one, two or three doses of the indicated vaccine or control. **B, C, D, E:** Different subclasses of TbpB-specific IgG in serum after one, two or three doses. 2-way ANOVA with Dunnett’s multiple comparison test comparing each TbpB immunized group to the unvaccinated control group used for calculating p value. **F:** Ratio of TbpB-specific IgG2b over IgG1 in serum after one, two or three doses. 2-way ANOVA with Tukey multiple comparison test comparing each TbpB adjuvanted group to all every other group used for calculating p value. **G, H:** Detection of anti-TbpB IgG and IgA in tracheal lavage (‘Nasal IgG/A’), and **I, J:** Detection of anti-TbpB IgG and IgA in vaginal lavages (‘Vaginal IgG/A’) after three doses of vaccine. Non-parametric Kruskal-Wallis test with Dunn’s multiple comparison test comparing each TbpB immunized group to the unvaccinated control group used for calculating p value. In all graphs, bars/lines, represent mean; error bars depict standard error. N = 6 per group (3 males depicted by squares; 3 females depicted by circles). Only significant comparisons shown on graph. ****, p<0.0001; ***, p<0.001; **, p<0.01; * p<0.05.

Further examination of IgG subclasses revealed that high levels of the Th2-associated IgG1 were elicited by all the vaccines after three doses (**Figure 1B**). Formulations containing Alum+MPL, Emulsigen D and CFA/IFA also produced robust levels of the Th1-associated IgG2b (**Figure 1C**), while only Emulsigen D and CFA/IFA elicited IgG2c to a significant extent (**Figure 1D**). Although typically associated with a response to polysaccharide antigens, anti-TbpB IgG3 was detected in groups that received Alum+MPL and Emulsigen D to a low, albeit significant, level (**Figure 1E**). The ratio of IgG2b/IgG1 was used as an indicator of Th1/Th2 bias (**Figure 1F**). After three doses, Alum and Addavax-formulations produced the lowest IgG2b/IgG1 ratio, indicative of a predominantly Th2 phenotype. Alum+MPL had a higher IgG2b/IgG1 ratio compared to Alum alone, but this difference was not statistically significant. Emulsigen D and CFA/IFA produced a significantly higher IgG2b/IgG1 ratio compared to Alum and Addavax, indicative of a balanced Th1/Th2 response. Although we had expected CFA/IFA to yield the higher levels of IgG2b compared to IgG1, in our hands, both subclasses were detected at similar levels.

Next, we quantified anti-TbpB IgG and IgA in mucosal lavages. Anti-TbpB IgG was detected, to varying extents, in nasal and vaginal lavages for all the adjuvanted vaccines, but only to a significant extent in vaginal lavages with Emulsigen D (**Figure 1G, I**). Anti-TbpB IgA was detected in some animals at low levels, but this was not significant (**Figure 1H, J**).

### Unique T cell cytokine responses induced by TbpB formulated with different adjuvants

To further assess cellular responses, splenocytes from naïve and vaccinated animals presented in Figure 1 were harvested approximately 2 weeks following the third dose, and then stimulated *ex vivo* with purified TbpB protein or media only. A T cell-focused 18-plex cytokine/chemokine array was conducted on the cellular supernatants in an effort to broadly detect antigen specific responses.

As predicted, naïve animals showed no detectable response in any analyte tested compared to media-stimulated control (**Figure 2 and <u>S1</u>**). Non-adjuvanted TbpB group demonstrated minimal immune response, comprised mainly of significantly elevated IL-5 compared to the naïve TbpB splenocyte control. TbpB+Alum formulation also had a minimal cytokine profile, comprising of significantly elevated levels of IL-4 compared to naïve TbpB control. Emulsigen D and CFA/IFA formulations had the most diverse immune profile eliciting a robust Th2 centric response (IL-4 and IL-5) while also producing the T regulator cytokine IL-10 and the Th17 cytokine IL-17A to a significantly higher level compared to naïve TbpB control. Alum+MPL and Addavax formulations also had unique immune signatures, comprised of all the previously mentioned cytokines except for IL-5 and IL-17A, respectively (**Figure 2A-E**).

**Figure 2:**
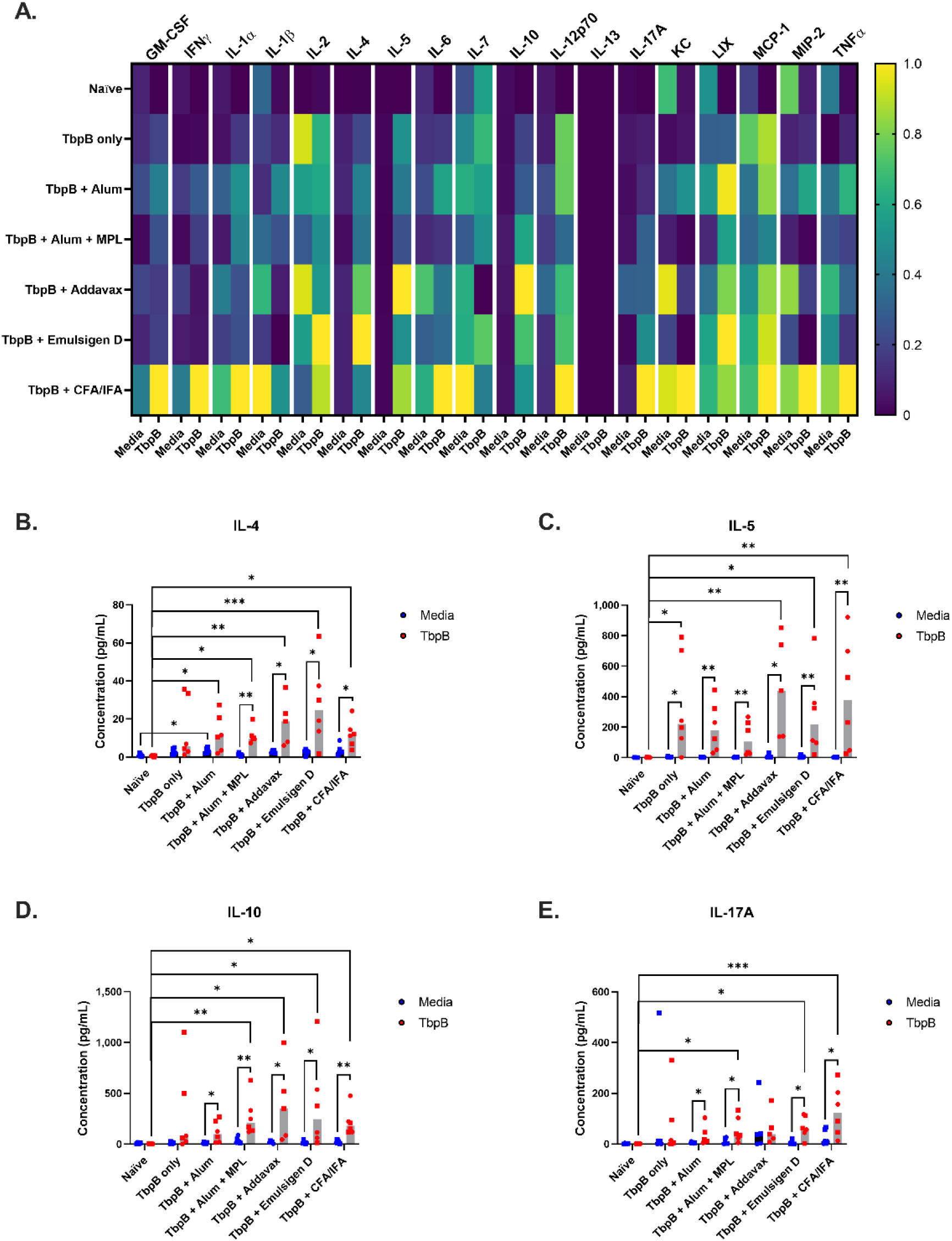
Multiplex cytokine Array from TbpB-vaccinated C57Bl/6 mice. Splenocytes from the indicated vaccine or control animals were stimulated with purified TbpB for 96 hours and supernatants were procured for cytokine and chemokine analysis. **A:** Median normalized heatmap. Median values for each cytokine were normalized to have the highest response as 1.0 and the lowest as 0 (n ≥ 5 mice per group). **B-E:** Data points represent an individual animal’s cytokine concentration to the indicated stimulator (media, blue; TbpB, red). Shaded gray bars represent the median and symbol shape indicates animal sex (males depicted by squares, females depicted by circles). Significance was determined using Dunn’s non-parametric multiple comparison with control for joint ranks and significant values are indicated on the graph. ***, p<0.001; **, p<0.01; * p<0.05. Additional graphed data and statistical analyses are shown in Figure S1.

Interestingly, despite being Th1 skewing in the literature, none of Alum+MPL, Emulsigen D or CFA/IFA demonstrated any vaccine-specific responses in IL-2, IFNγ or TNFα at the time point examined (**<u>Figure S1B, E, N</u>**). There were also no other trends of note in any other cytokine/chemokine tested (**Figure S1**). These results indicate that each of these adjuvants paired with the same protein antigen elicit a unique T cell signature.

Taken together with the previous figure, we confirmed that different adjuvants in conjunction with the same antigen led to distinctive humoral and cytokine signatures, prompting us to further investigate whether there is an impact on mucosal clearance of pathogenic Neisseria.

### Adjuvants impact vaccine-mediated clearance of N. gonorrhoeae from the lower genital tract of female mice

To account for any inherent differences arising due to the genetic background of inbred mice, we utilized both wild type C57BL/6 and BALB/c females to evaluate protection against lower genital tract infection by *N. gonorrhoeae.* Three doses of Ngo MS11 TbpB vaccine formulated with five different adjuvants were administered 21 days apart. Prior to challenge, mice that had entered the diestrus stage of the estrus cycle were treated with β-estradiol to arrest the reproductive cycle and antibiotics to suppress commensals. Mice were then inoculated vaginally with *N. gonorrhoeae* MS11 and daily vaginal lavages were collected to enumerate bacterial burden and track the duration of colonization. Protection was assessed in two independent experiments for each vaccine formulation in the C57BL/6 background, and a single study in the BALB/c background (See **<u>Table S1</u>**).

Overall, the median duration of colonization in all vaccinated cohorts was reduced compared to naïve controls in both mouse backgrounds (~3-5 days in vaccinated versus ~8 days in naïve for C57BL/6 mice; ~6-8 days in vaccinated versus 14 days in naïve for BALB/c mice) (**Figure 3A-E, G-K**). However, only Alum+MPL and Emulsigen D formulations gave a reduction in % colonization over time that was statistically significant compared to naïve controls in the BALB/c background (p=0.0222 and p=0.0355, respectively) and only the Emulsigen D cohort approached statistical significance (p=0.0562) in the C57BL/6 background. Bacterial burden in individual animals was highly variable from day to day (**<u>Figure S2</u>**), which has previously been reported in this model [31]. So, despite the accelerated clearance, there was no significant difference in the total bioburden, estimated using area under the curve (**Figure 3F, L**).

**Figure 3:**
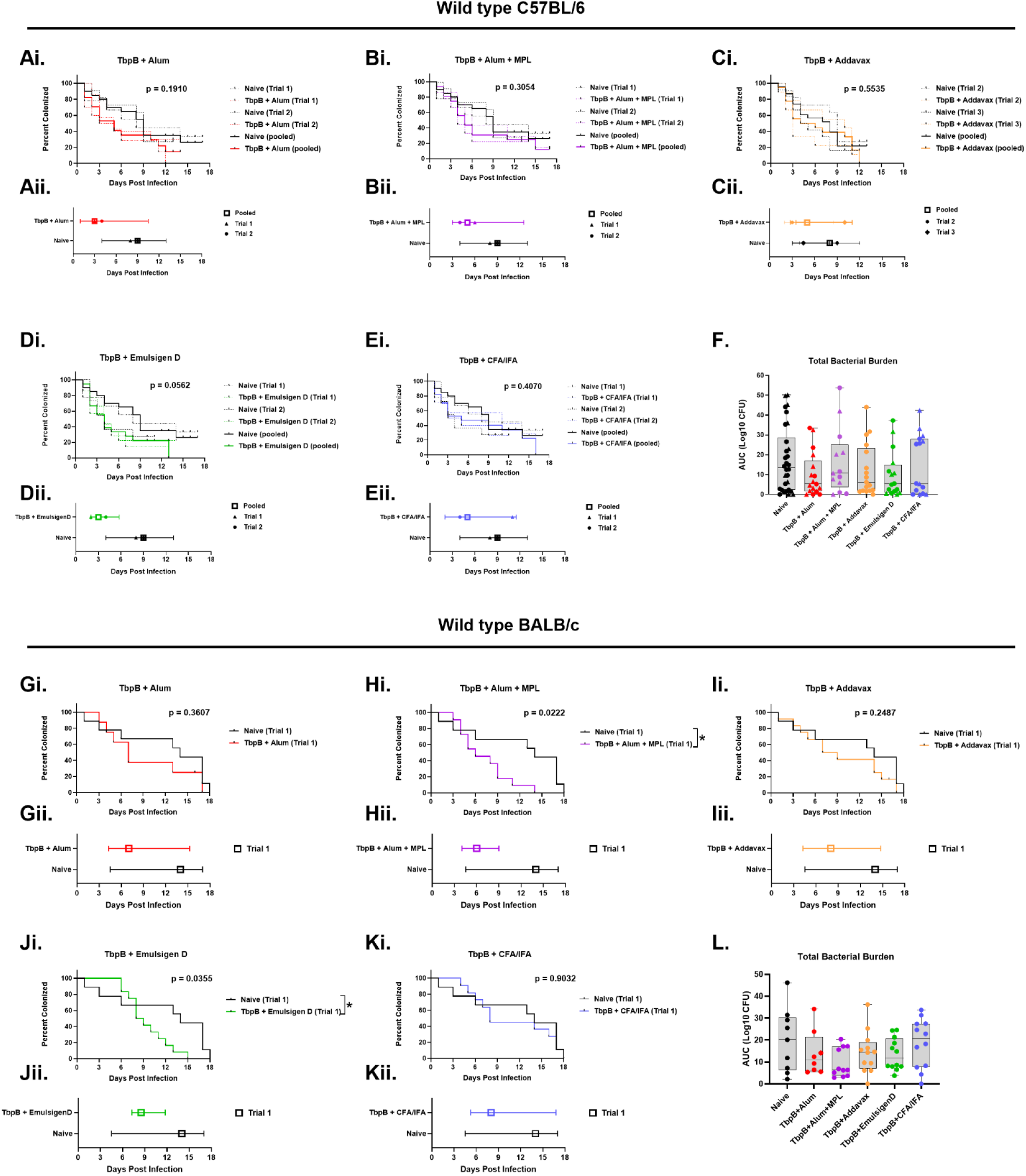
Protection conferred by different TbpB-based formulations against gonococcal colonization in the female lower genital tract. **A-E, G-K (panel i):** Percentage of wild type C57BL/6 or BALB/c females that remain colonized in the lower genital tract after gonococcal infection. Each curve compares naïve (black line, same group for each panel) controls to 3x TbpB-vaccinated (coloured line, group specified) formulations indicated above. All groups were evaluated together but have been separated into individual panels for ease of interpretation. p-value calculated using Log-Rank (Mantel-Cox) curve comparison test indicated on each graph. *, p<0.05. **A-E, G-K (panel ii):** depicts the median colonization duration (day on which 50% of the cohort cleared infection) and interquartile range. **F, L:** Box-plots showing total bacterial burden in each animal estimated by the area under the curve of log-transformed daily CFU recovery data in Figure S2. p-values were obtained using non-parametric Kruskal-Walllis test with Dunn’s multiple comparison were non-significant. N=8-12 (BALB/c), N=13-19 (pooled independent C57BL/6 trials), see Table S1 for detailed breakdown. For C57BL/6, Trial 1 and Trial 2 data pooled for TbpB + Alum, TbpB + Alum + MPL, TbpB + Emulsigen D, TbpB + CFA/IFA comparisons to Naïve in A,B,D,E; Trial 2 and Trial 3 data pooled for TbpB + Addavax comparison to Naïve in C; All animals included in F.

These studies demonstrate adjuvants impact vaccine-mediated clearance of gonococci from the lower genital tract of mice, with an increased protective effect observed when using two supposed Th1/Th2-balanced adjuvants based on literature, Alum+MPL and Emulsigen D.

### Adjuvants influence vaccine-mediated clearance of N. meningitidis from the nasopharynx of humanized transgenic mice

To compare the efficacy of different adjuvants in a *N. meningitidis* colonization model, human CEACAM1 (hCEACAM1) transgenic mice were immunized three times with formulations of Nme B16B6 TbpB vaccine followed by intranasal infection with the matched Group B strain B16B6, and sacrificed 3 days post infection to enumerate bacterial burden in the nasopharynx. Transgenic mice were utilized in these studies since hCEACAM1 serves as the receptor for Neisserial Opa adhesins and is essential for nasopharyngeal colonization in mice [32]. Protection was assessed in two independent experiments in each of the two separate CEACAM1-humanized genetic backgrounds, C57BL/6 and FvB, to ensure reproducibility.

Bacterial burden in the nasopharynx, pooled from the two independent experiments for each mouse line, is depicted (**Figure 4Ai and 4Bi**). Compared to naïve controls, a reduction in the median bacterial burden was noted for all the immunized groups, with Alum+MPL and CFA/IFA producing a significant drop in recovered colony forming units (CFUs) in both C57BL/6 and FvB backgrounds, and Addavax leading to a statistically significant outcome in the FvB background and approaching significance in C57BL/6.

**Figure 4:**
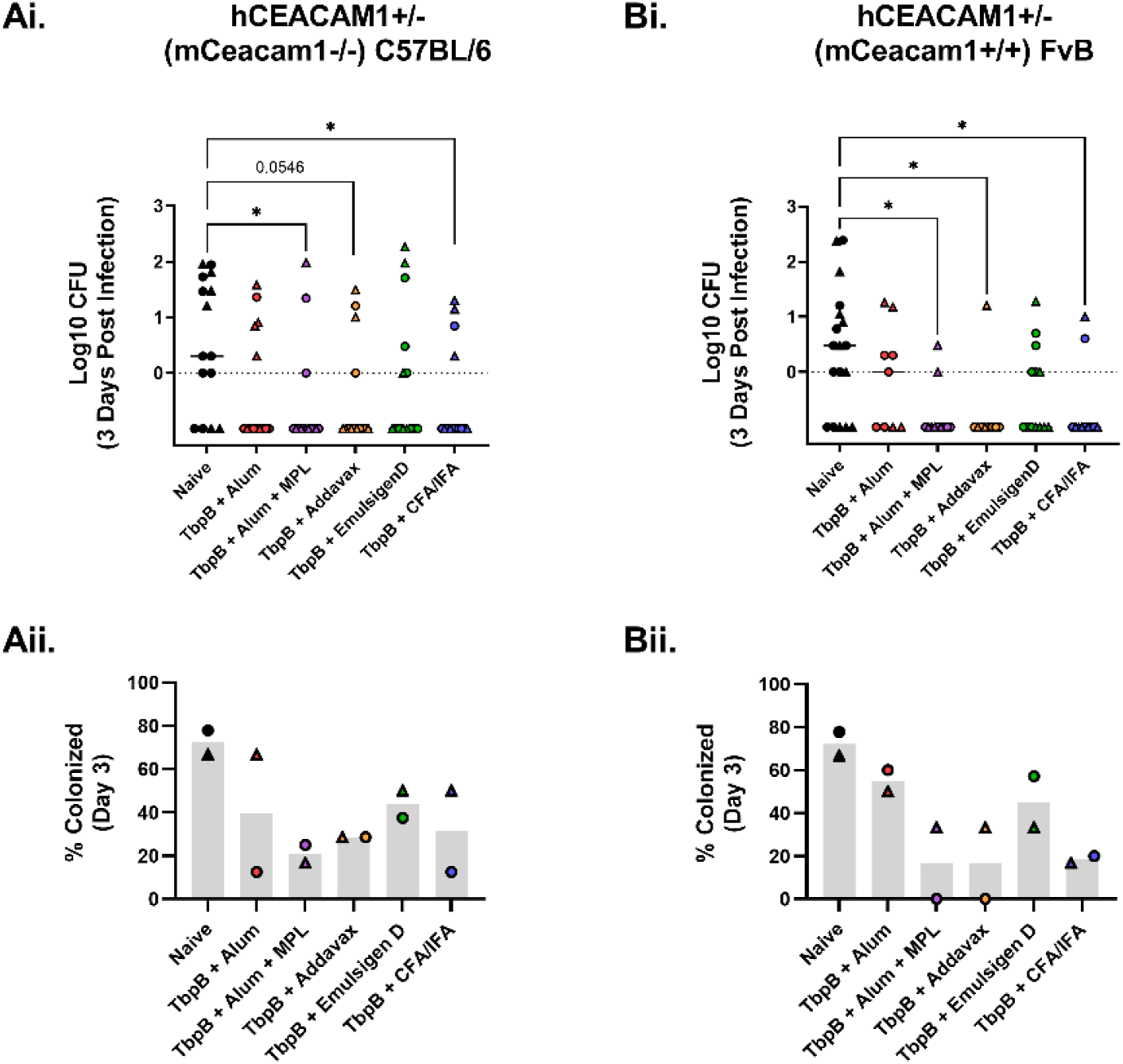
Protection conferred by different TbpB-based formulations against meningococcal colonization in the upper respiratory tract. **Ai, Bi:** Bacterial recovery 3 days post infection from the nasopharynx of naïve or three times TbpB-vaccinated transgenic hCEACAM1+/− (mCeacam1−/−) C57BL/6 and hCEACAM1+/− (mCeacam1+/+) FvB mice. Data from individual animals from two independent experiments, indicated by either circle (Trial 1) or triangles (Trial 2), depicted. N (pooled) = 9-18 per group; detailed breakdown provided in Table S1. Line depicts median. Non-parametric Kruskal-Wallis test with Dunn’s multiple comparison was performed to compare each TbpB immunized group to the unvaccinated control group. Only significant or close to significant p-values shown. * p<0.05. **Aii, Bii:** Percentage of culture positive animals 3 days post infection in the two different transgenic lines indicated above. Bar depicts median. Percent colonized in two independent experiments, indicated by either circle (Trial 1) or triangles (Trial 2).

The percentage of culture positive animals was overall lower in vaccinated mice compared to naïve controls (68% and 78% colonized in each independent C57BL/6 and FvB trial, respectively) (**Figure 4Aii, 4Bii**). The outcome was most prominent in Alum+MPL (25% and 17%, 0% and 33% colonized in C57BL/6 and FvB, respectively), CFA/IFA (13% and 50%, 17% and 20% colonized in C57BL/6 and FvB, respectively), and Addavax (29% and 29%, 0% and 33% colonized in C57BL/6 and FvB, respectively). Alum (12.5% and 67%, 50% and 60% colonized in C57BL/6 and FvB, respectively) and Emulsigen D (38% and 50%, 33% and 58% colonized in C57BL/6 and FvB, respectively), led to a modest reduction and/or were more variable across replicates.

Similar to our observations in the genital gonococcal model, vaccine efficacy against *N. meningitidis* in the nasopharynx is also impacted by the choice of adjuvant. Intriguingly, the best performers included adjuvants Addavax, Alum+MPL and CFA/IFA, which are all vastly distinct formulations with distinct immune and functional characteristics.

### No consistent link between antibody levels and gonococcal clearance from the lower genital tract

While the effector processes that confer mucosal protection remain elusive, antibodies have been implicated in protection against both genital gonococcal and nasopharyngeal meningococcal infections in mouse models [35, 37]. To investigate whether a correlation exists between antibody levels and protection, total IgG, IgG1 and IgG2b in pre-challenge or terminal serum, as well as IgG and IgA levels in appropriate mucosal secretions, were quantified in animals used in challenge studies and the data segregated by colonization status (**Figure 5** and **Figure 6**).

**Figure 5:**
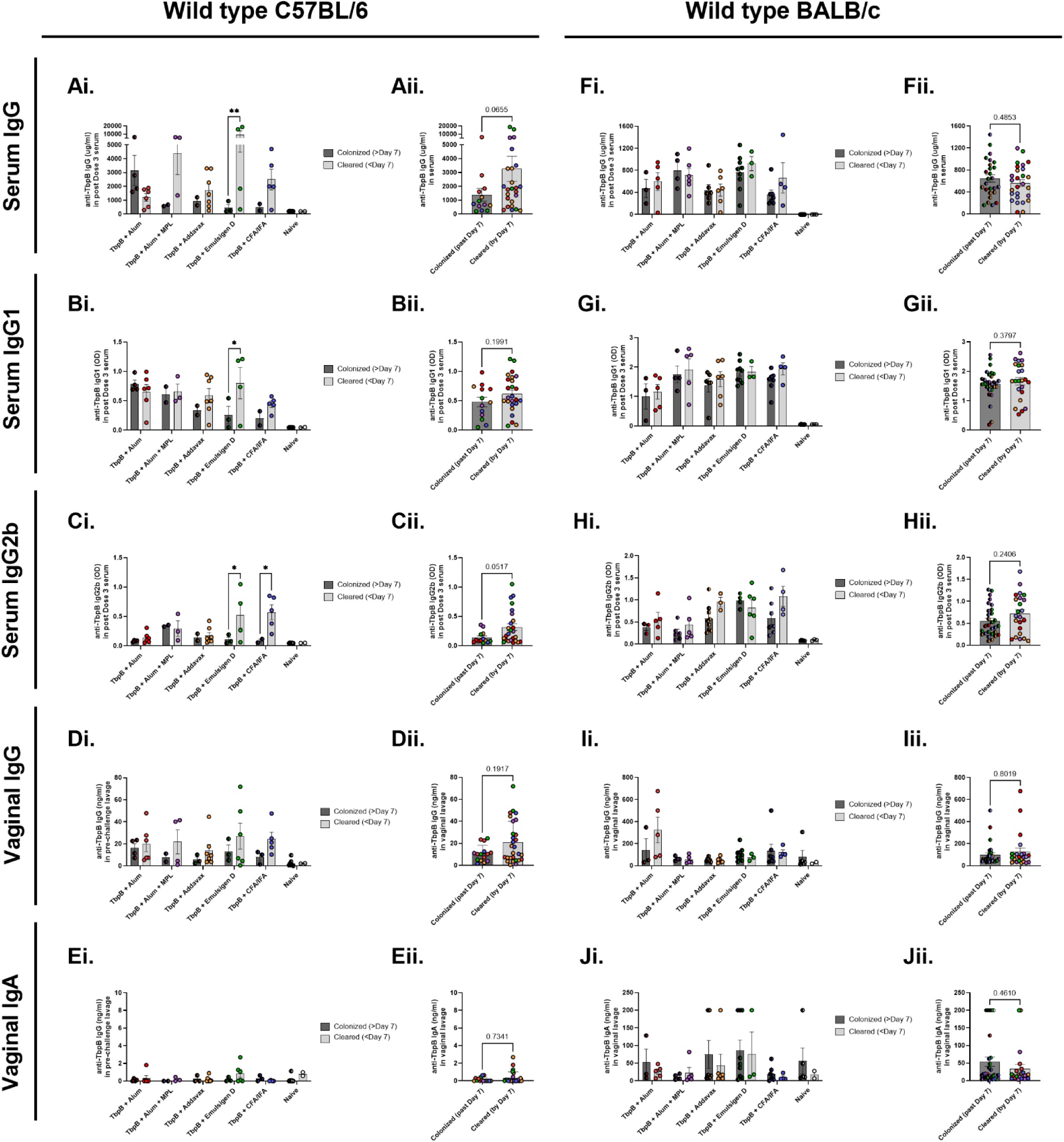
Comparison of *α*-TbpB antibody levels in animals that cleared gonococcal infection within 7 days versus those that remained colonized past 7 days. **A, F:** Total serum IgG; **B, G:** Serum IgG1; **C, H:** Serum IgG2b; **D, I:** Vaginal IgG; **E, J:** Vaginal IgA from Ngo challenge studies segregated by colonization status at 7 days post infection. Panel i provides further breakdown within each vaccine or control group; Panel ii, segregated by colonization status for all TbpB-vaccinated animals combined. Bars represent mean; error bars depict standard error. N=6-10 per group for C67BL/6, Trial 2 samples. N=8-12 per group for BALB/c; Trial 1 samples. detailed breakdown found in Table S1. For Panel i, 2-way ANOVA with Sidak’s multiple comparison test was used for calculating p-values; only significant comparisons displayed. **, p<0.01; *, p<0.05. For Panel ii, non-parametric, two-tailed, Mann-Whitney test was used, and p-values indicated on each graph.

**Figure 6:**
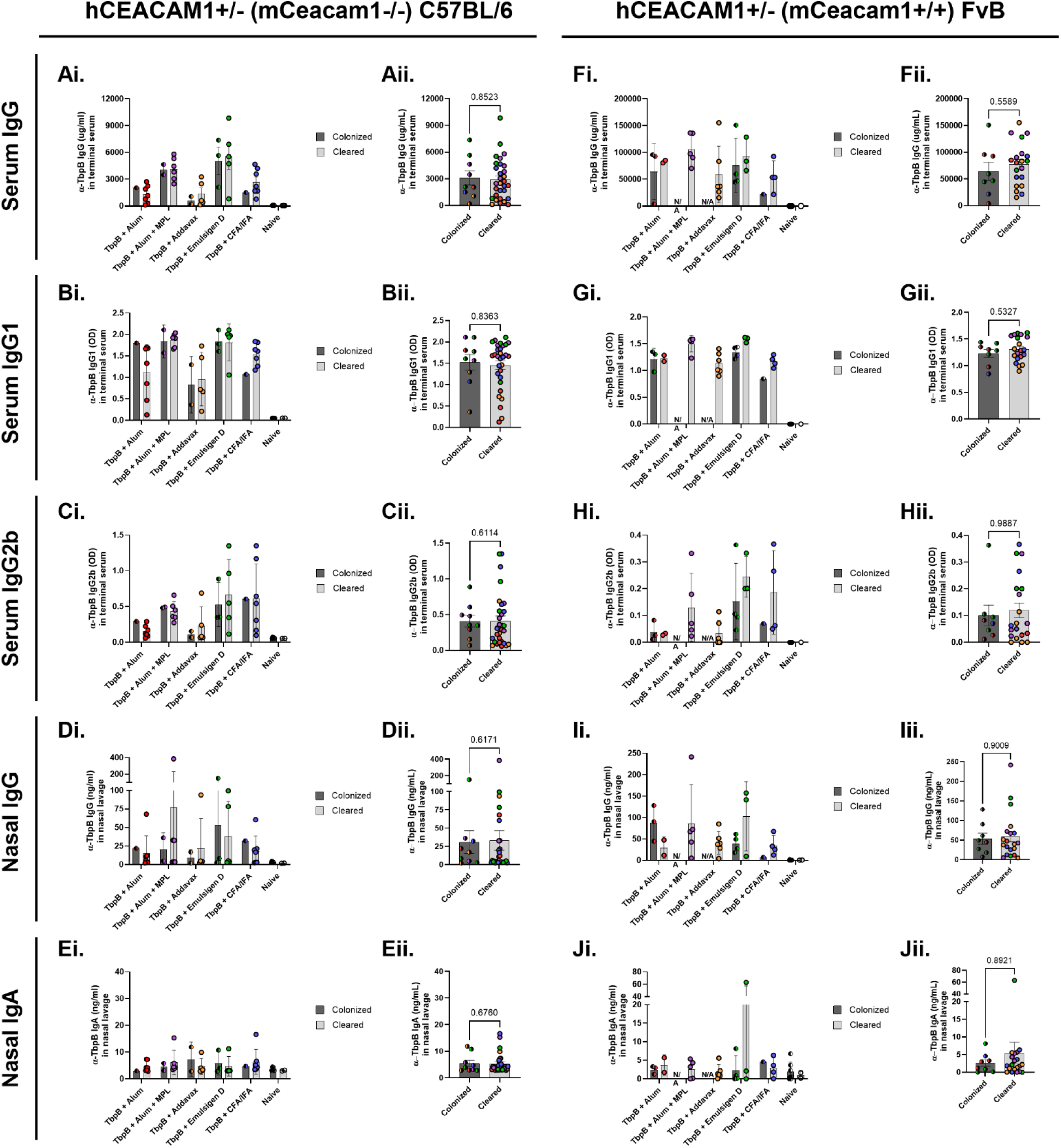
Comparison of *α*-TbpB antibody levels in animals that cleared meningococcal infection within 3 days versus those that remained culture positive at 3 days. **A, F**: Total serum IgG; **B, G:** Serum IgG1; **C, H:** Serum IgG2b; **D, I:** Nasal IgG; **E, J:** Nasal IgA from Nme challenge studies segregated by colonization status at 3 days post infection. Panel i provides further breakdown within each vaccine or control group; Panel ii, segregated by colonization status for all TbpB-vaccinated animals combined. Bars represent mean; error bars depict standard error. N=7-9 per group for hCEACAM1+/− (mCeacam1−/−) C67BL/6, N=5-8 per group hCEACAM1+/− (mCeacam1+/+) FvB; Trial 1 samples analyzed for both transgenic lines; detailed breakdown of N found in Table S1. For Panel i, 2-way ANOVA with Sidak’s multiple comparison test was used for statistical analysis; non-significant p-values not displayed. For Panel ii, non-parametric, two-tailed, Mann-Whitney test was used, and p-values indicated on each graph.

Since all animals eventually clear infection in the gonococcal model, we used colonization status on Day 7 post infection to assess whether there were differences in antibody levels in animals that cleared infection “early” (<Day 7) versus “late” (>Day 7). Curiously, Alum+MPL and EmulsigenD, the two groups in which accelerated gonococcal clearance was significant (**Figure 3**), had the highest overall serum IgG (**Figure 1**, **<u>Figure S3A, F, K, P</u>**) making it enticing to consider whether animals with higher antibody titres in general cleared infection “early”.

Segregating by colonization and vaccine status revealed significantly higher serum IgG, IgG1, IgG2b only in Emulsigen D-formulated and higher serum IgG2b in CFA/IFA-formulated C57BL/6 mice that cleared infection “early” (**Figure 5Ai, Bi, Ci**). However, this difference was not seen for the other vaccine formulations in the C57BL/6 background. IgG (p=0.0655) and IgG2b (0.0517) levels appeared elevated after pooled “early” versus “late” animals from all the immunized C57BL/6 mice, but the difference was not statistically significant (**Figure 5Aii, Cii**). Interestingly, in the BALB/c background there was no difference in serum IgG, IgG1, IgG2b (**Figure 5Fi, Gi, Hi, F-H, <u>panel ii</u>**), signifying that the trends observed with certain groups in C57BL/6 cannot be generalized. We also found no difference in vaginal IgG or IgA levels at baseline and colonization status in either background (**Figure 5D, E, I, J**).

### No correlation between antibody levels and meningococcal clearance from the nasopharynx

Considering that two of the most protective formulations in this model, with adjuvants Alum+MPL and Addavax, had some of the highest and lowest serum IgG, IgG1, IgG2b titres respectively, serum antibody levels may not be a good predictor of nasopharyngeal protection. Nevertheless, we performed a similar pairwise analysis for the meningococcal challenge samples. Since bacterial enumeration in the nasopharynx on Day 3 post infection is a terminal procedure, we compared animals that had cleared infection versus remained culture positive on Day 3.

Indeed, pairwise comparison showed no difference within each group or when cleared versus colonized animals across the immunized groups were pooled (**Figure 6A-C, F-H**). There was also no difference in IgG and IgA levels in nasopharyngeal lavages collected on Day 3 (**Figure 6D, E, I, J**). The lack of any correlation was consistent in both the C57BL/6 and FvB transgenic animals.

In conclusion, we did not find a generalizable correlation between immunogen-specific antibody levels present in serum or mucosa and accelerated mucosal clearance of either gonococci or meningococci in the mouse models utilized here.

### No evident correlation between SBA titre and Neisserial clearance

SBA has long been used as a correlate of protection against invasive meningococcal disease, prompting us to examine whether it can be used as a correlate for mucosal protection. For the gonococcal studies, heat inactivated terminal serum that was collected approximately 21 days post-challenge was used in conjunction with pooled human serum against the challenge strain MS11 in SBA assays. Bacteria was iron starved to emulate iron restriction encountered during infections, which induces transferrin receptor expression. It is worth noting that naïve infected animals did not produce a SBA titre, signifying that any bactericidal activity observed in the immunized groups can be attributed to vaccination. However, only a subset of immune serum was found to be bactericidal; Alum+MPL formulation gave the highest proportion of samples that were bactericidal in both C57BL/6 and BALB/c mice (**Figure 7Ai, Bi**). Pairwise comparison of animals that cleared infection “early” (<Day 7) versus “late” (>Day 7) within each group did not reveal any significant trends in SBA titres (**Figure 7Aii, Bii**), nor did pooling all “early” and “late” immunized animals (**Figure 7Aiii, Biii**).

**Figure 7:**
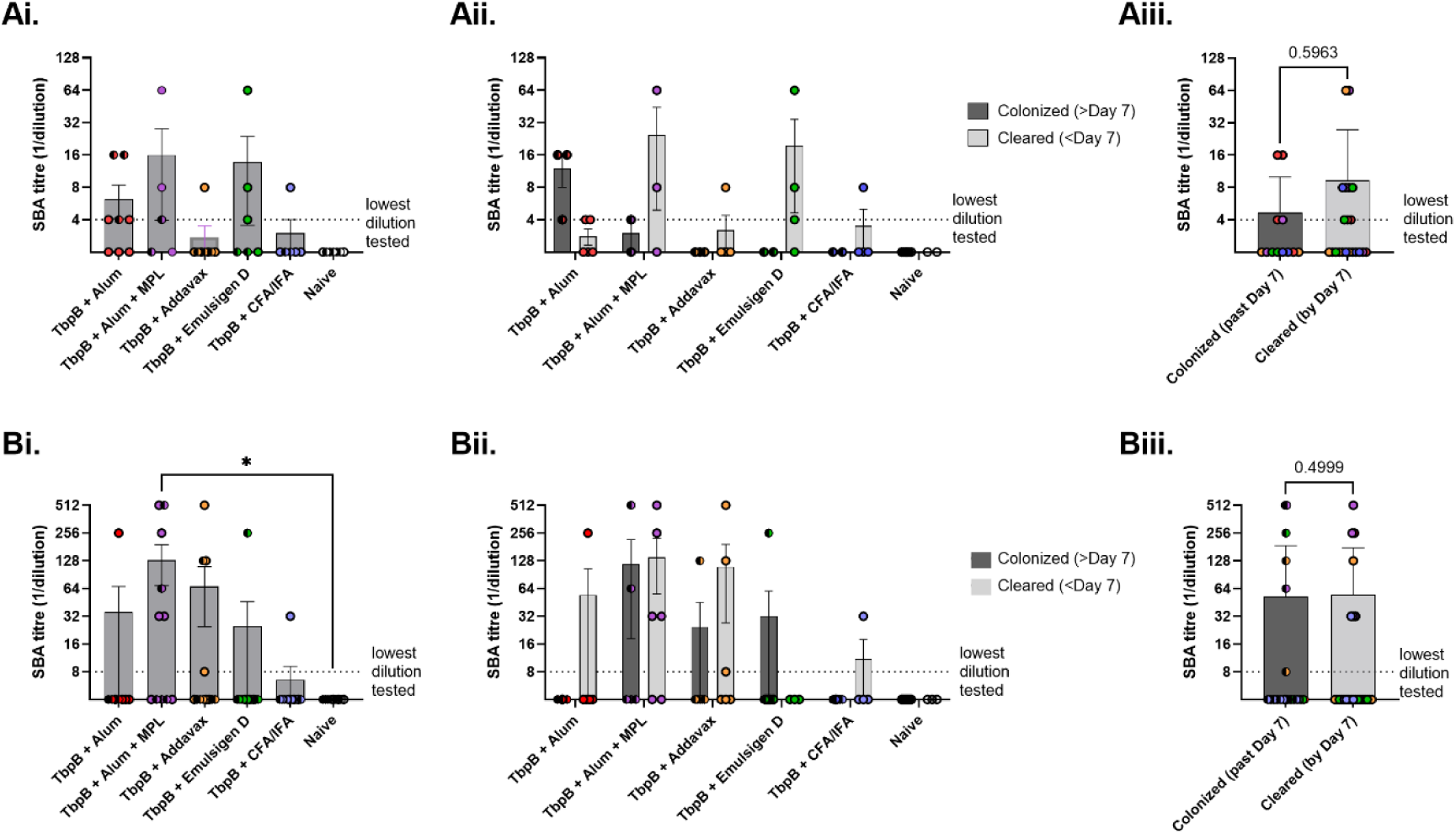
Comparison of SBA tires in animals that cleared gonococcal infection within 7 days versus those that remained colonized past 7 days. **Ai, Bi:** SBA titres against iron-starved Ngo MS11 strain elicited by different TbpB-based vaccines in C57BL/6 and BALB/c mice, respectively. Non-parametric Kruskal-Wallis test with Dunn’s multiple comparison used for calculating p-value. *, p<0.05; only significant p-values displayed. **Aii, Bii:** SBA titres segregated by vaccine received and colonization status at 7 days post infection in C57BL/6 and BALB/c mice, respectively. 2-way ANOVA with Sidak’s multiple comparison test was used for calculating p-values; non-significant p-values not displayed. **Aiii, Biii:** SBA titres segregated by colonization status for all TbpB-vaccinated animals combined. non-parametric, two-tailed, Mann-Whitney test was used, and p-values indicated on each graph. N=5-8 per group for C57BL/6, Trial 2 samples. N=8-12 per group for BALB/c; Trial 1 samples. Detailed breakdown found in Table S1.

Surprisingly, despite observing protection (**Figure 4**) in the meningococcal model, heat inactivated terminal sera from matched animals were not bactericidal against the challenge strain B16B6 using either baby rabbit complement or pooled human serum as the complement source (data not shown). Overall, with the limited number of serum samples for which SBA titre was achieved, we observed no definitive correlation between SBA titre and mucosal protection.

## DISCUSSION

Although current vaccine development efforts are directed primarily towards Ngo, it is important to recognize that attaining robust pan-Neisserial protection through a universally effective vaccine holds significant potential to enhance overall vaccine adoption, particularly given that the general acceptance for widespread implementation of a gonococcal vaccine may face significant barriers, including low self-perceived risk of contracting sexually transmitted infections (STI) among the general population, parental attitudues towards the implication of a STI vaccine on adolescent sexual behaviour among several factors (reviewed in [50]). Additionally, while Ngo mainly occupies the genital tract and Nme the nasopharynx, extra-genital infections by Ngo are frequent and oropharyngeal infections act as a major contributor to transmission (reviewed in [51]), while the recent identification of Nme urethritis cases has led to the identification of a urethral clade that has emerged from Nme serogroup C strains [52]. Therefore, developing Neisserial vaccines that are effective at multiple mucosal sites will be vital to ongoing vaccine efficacy. While the selection of appropriate vaccine antigen(s) is critical during early development, understanding the optimal type of immunological response needed for protection and fine-tuning the adjuvant component to achieve this desired response is equally vital.

To our knowledge, this is the only comparative study to examine vaccine-driven protection using both *N. gonorrhoeae* and *N. meningitidis* colonization models. We have used TbpB as an immunogen due to the well-established role of the transferrin receptor in facilitating bacterial survival during both gonococcal and meningococcal infections, as well as its demonstrated utility as a vaccine antigen [25, 53–56]. Despite its inherent sequence variability, a challenge addressed in this study by immunizing with the matched TbpB from the challenge strains, we have recently demonstrated the feasibility of achieving broad vaccine coverage by incorporating multiple TbpB variants into a single formulation [57]. Nevertheless, while the focus was on TbpB in this study, we anticipate that the insights gained will hold broader relevance to other soluble protein antigens as well.

The goal of the adjuvant panel selected here was to encompass a spectrum of biochemical and immunological differences. At the same time, we intentionally narrowed our choices to include adjuvant components used in commercial vaccines, adding relevance and practicality to our findings. In our hands, the adjuvants showed evidence of Th-biases reported in literature [41–49]. A limitation of our immunological characterization is that we had to rely upon IgG subclasses for an indication of Th bias. Although our cytokine panel was able to provide a high-level assessment of cytokine signatures associated with each of the formulations, it was unable to resolve antigen-specific Th1 responses. While direct visualization by flow cytometry would have been an ideal approach, we were unable to determine antigen-specific T cell responses in splenocytes by flow cytometry. This was despite significant efforts, including the use of either purified TbpB or a peptide pool as the stimulant. Future work must focus on the development of neisserial antigen-specific T cell assays to overcome this limitation. For the scope of this study however, the observation of differentiating humoral and cytokine responses provided sufficient confidence to proceed to challenge phase.

Differences in vaccine performance was nuanced with accelerated bacterial clearance observed across all five formulations in both models. In the gonococcal model, Alum+MPL and Emulsigen D were the best performing adjuvants. While the commonality between these two may be their ability to elicit a Th1/2 balanced response, it is important to keep in mind that each adjuvant activates different pathways at the time of immunization. MPL is a TLR4 agonist that directly activates antigen presenting cells and leads to an inflammatory environment favouring Th1 differentiation [42, 43]. Thus MPL, in conjunction with the Th2-skewing Alum, should confer a balanced response. Emulsigen D, an oil-in-water emulsion adjuvant used in veterinary vaccines is less well characterized. It contains DDA, which form cationic liposomes that associate with antigen and enhance uptake by antigen presenting cells [48]. Unexpected in considering that it tends to elicit a strong inflammatory response, CFA/IFA did not perform as well in the gonococcal model. One key difference between CFA/IFA and other adjuvanted formulations was that CFA/IFA-formulations were administered subcutaneously due to its potent reactogenicity, while the rest were provided intraperitoneally. Intraperitoneal immunization has been associated with faster antigen drainage to lymphoid organs and more robust CD8 activation [58]; however, the relevance of these functions for gonococcal clearance is not clear. Following up on different routes of immunization and its impact on gonococcal protection is a fascinating avenue for future endeavours and will provide more relevant information for human vaccines going forward.

For *N. meningitidis*, the results were more perplexing. Robust performance was observed with Th2-skewing Addavax but not Alum, and Th1/2-balanced Alum+MPL and CFA/IFA but not Emulsigen D. These observations indicate the need for a more comprehensive assessment of immunological factors to deduce common features important for protection in the upper respiratory tract. Interestingly, Alum+MPL was the common adjuvant that gave robust protection in both models. Clinical-grade MPL is a constituent in the commercial human papillomavirus vaccine (Cervarix®) in conjunction with Alum, as well as in shingles and malaria vaccines (Shingrix® and Mosquirix®, respectively) as liposome-based formulations with the added immunostimulant QS-21 [59]. The widespread acceptance and safety profile of MPL-containing vaccines in recent years makes the inclusion of MPL or its synthetic analogue, glucopyranosyl lipid (GLA) [47, 60] a practical consideration for novel neisserial vaccines.

A key consideration in our study was the use of two genetically and phenotypically distinct mouse lines to ensure that our observations were reproducible, regardless of inherent immunological differences in inbred mice that could potentially impact clearance mechanisms. For instance, lower genital tract gonococcal infection in BALB/c mice is inflammatory, with a spike in pro-inflammatory cytokines and a concomitant influx of neutrophils occurring approximately 5 days post infection, whereas a proinflammatory response is not typically seen in C57BL/6 mice [61, 62]. On the other hand, meningococcal clearance in FvB mice has to be independent of complement mediated lysis due to a defect in the C5 gene, which is intact in C57BL/6 [63]. Despite these differences, performance of the various formulations was consistent across mouse backgrounds. A key limitation of our study was the reliance on naïve mice as the sole control group. Although adjuvant controls would have provided a more rigorous comparison, the inclusion of five additional groups posed ethical and practical challenges given the scale of the experiments. Another limitation is the sex-restriction of the gonococcal model to female animals only. While efforts are underway to establish a gonococcal urethritis model in male mice and a gonococcal nasopharyngeal colonization model that can utilize both males and females, these new models have not yet been validated for vaccine studies. It is worth noting that no sex dependent effects have been observed in the initial non-challenge study or in the meningococcal challenge studies where both sexes were included.

We chose to focus on humoral responses to study correlates of protection due to evidence of anti-LOS antibody mediated gonococcal clearance in mice and the established correlation between bactericidal titres and protection against invasive meningococcal disease [8, 35]. In the gonococcal model, bacterial clearance over time even in naïve animals is expected, due to several factors that vary from animal to animal. These include basal inflammation and complex interplay between Ngo and vaginal microbiome potentially impacting initial colonization rates, pro-inflammatory responses and influx of PMNs starting approximately 5 days post infection in BALB/c mice [61], the continuation of natural cycling around 9 days post-infection [62], and the overgrowth of commensals leading to Ngo being outcompeted and contaminants appearing on plates after approximately 1 to 2 weeks. Therefore, Day 7 was used as a cutoff for differentiating “early” versus “late” clearance, when ~50-70% of mice across the various vaccinated groups had cleared infection compared to ~30% in the controls, as it allowed for appropriate sample distribution to draw comparisons while avoiding confounding effects due to the reasons outlined. For the meningococcal model, we were restricted to using Day 3 as the cutoff due since bacterial enumeration using this model is a terminal procedure. We were unable to find a correlation between antibody titres and protection in this study, and this is consistent with independent findings by our group when using other vaccines. However, it needs to be emphasized that a lack of correlation does not diminish any potential role played by antibodies during bacterial clearance. Monitoring antibody titre remains a vital metric for assessing the development of a robust immune response post vaccination. Our data underscores the importance of interpreting absolute titres carefully, cautioning against dismissing a formulation solely based on lower antibody titres. This reinforces the importance of relevant infection models to understand vaccine-mediated protection against these or other pathogens.

Based on our observations, we conclude that beyond measuring antibody titres, a thorough cellular analysis of affected tissues and lymphoid organs at various time points post-infection would likely be necessary to assemble a comprehensive dataset for multivariate analysis. Since the aim of this study was to broadly survey the efficacy of different adjuvant formulations with TbpB, such detailed analysis of cellular responses was beyond our scope. Forthcoming adjuvant-controlled studies should now focus on utilizing a single formulation to delve deeper into the mechanisms underlying bacterial clearance and, ideally, reveal a definitive correlate of protection.

In summary, this study underscores the importance of adjuvant optimization in achieving optimal efficacy and highlights the limited predictive ability of antibody titres alone in assessing vaccine-mediated protection. Moreover, it reinforces that protection against infection of one mucosal site does not necessarily predict protection at other sites, but that judicious choice of adjuvant will allow a single vaccine to confer protection at both sites. These findings provide guidance for the design and evaluation of novel formulations with these and other immunogens, ultimately advancing our collective efforts in combatting neisserial pathogens.

## METHODS

### Protein purification

Protein expression vector was heat-shocked into chemical-competent E.coli T7express cells (New England Biolabs) and allowed to recover in LB broth for 1 hour, shaking at 175 RPM and 37°C, before selection by additional LB broth (3 mL) supplemented with ampicillin (100 μg/mL) and grown for a further 3 hours. The starter culture was used to inoculate 6 L of autoinduction media [64] with 50 μg/mL of ampicillin. The culture flask was then shaken overnight at 37°C and 175 RPM for 16 hours and a further 24 hours at 20°C. Cells were harvested by centrifugation at 5000 x g’s for 30 minutes at 4°C, resuspended in 50 mL of resuspension buffer (50 mM Tris pH 8.0, 300 mM NaCl) supplemented with protease inhibitors (cOmpleteTM Mini, Roche), 1 mg/mL lysozyme, and 0.03 m/mL DNase I. Cells were homogenized (EmulsiFlex-C3, Avestin) and the resulting lysate was clarified by centriguation at 16,000 x g’s for 90 minutes at 4°C, followed by 0.22 μm syringe filtration (Millipore). The clarified lysate is then recirculated by a peristaltic pump and affinity captured on a HisTrap EXCEL FF column (Cytiva). The column is then mounted onto an ÄKTA Purifier 100 (GE Healthcare), washed and then eluted with imidazole (50 mM Tris pH 8.0, 300 mM NaCl, 300 mM imidazole pH 7.4). Eluted fraction was dialyzed overnight into exchange buffer (50 mM Tris pH 8.0, 150 mM NaCl) and sample purity is assessed by 12% SDS-PAGE analysis and protein concentration determined by Nanodrop (ThermoFisher) absorbance at 280 nM. In-house purified TEV protease is added to separate the N-terminal MBP fusion partner from TbpB and the sample was then exchanged into a low-salt buffer (50 mM Tris pH 8.0, 10 mM NaCl) and loaded onto an anion exchange HiTrapQ FF column (Cytiva) and the fraction containing TbpB was concentrated and passed through a size exclusion column (HiPrep 26/60 Sephacryl S-200 HR, Cytiva) to further remove any contaminants and exchange into the final sample buffer (50 mM HEPES, 50 mM NaCl). A final polishing step with strong anion exchange MonoQ 5/50 GL column (Cytiva) to remove lipopolysacchrides and the final sample was concentrated and aliquots flash frozen by liquid nitrogen. Final edotoxin content was quantified using Limulus Amebocyte Lysate (LAL) assay kit (Waki Chemicals, cat. WPEK4-20015 according to manufacturer’s instructions.

### Vaccine preparation and immunizations

Vaccine formulations were prepared with either purified MS11 TbpB or B16B6 TbpB protein in phosphate buffered saline (PBS) (<0.05 EU endotoxin per µg protein). Alum: antigen was first diluted in PBS, 2% Alhydrogel (Sigma-Aldrich) was then added to give a final concentration of 25 μg protein and 7.7% (v/v) 2% alhydrogel per 100 μL dose and allowed to adsorb for at least 20 mins with gentle agitation. Alum+MPL: same preparation as Alum, except 10 µg MPLA in DMSO (Invivogen, cat. Tlrl-mpla2) was added as a last step resulting in a final concentration of 25 μg protein, 7.7% (v/v) 2% alhydrogel, 10 μg MPL in 10% (v/v) DMSO per 100 µL dose. Addavax (Invivogen, cat. Vac-adx-10) and Emulsigen D (MVP Adjuvants): ready to use emulsions were added to diluted protein and gently mixed to prepare 25 μg protein, 50% (v/v) Addavax or 20% (v/v) Emulsigen D per 100 µL dose. CFA/IFA (Invivogen. Cat. vac-cfa-10, vac-ifa-10): adjuvant was added to diluted protein at a 60% (v/v) ratio, mixed by pipetting several times, then homogenized for 3 x 15 seconds at ~8,000 rpm using IKA handheld homogenizer to form a stable emulsion. All formulations were injected intraperitoneally, with the exception of CFA/IFA which was subcutaneously administered. Saphenous bleeds were collected ~2 weeks after each dose to track the antibody response.

### *Ex vivo* mouse splenocyte stimulation and supernatant cytokine/chemokine array

4-6 week old C57BL/6 mice (purchased from Charles River) were allowed to acclimatize for a week and then vaccinated with 25 ug *Nme* B16B6 TbpB with indicated adjuvant intraperitoneally [except CFA/IFA samples, which were subcutaneously administered] three times with 3 weeks intervals, and spleens were harvested two weeks after the last vaccination. Splenocytes were isolated by pressing spleens through a 40 μm cell strainer to remove tissue debris and then treated with red blood cell lysis buffer (eBiosciences. cat. 00-4300-54). Cells were suspended (1 × 10^6^ cells per well) in complete RPMI medium (cRPMI; RPMI 1640 supplemented with 10% heat-inactivated FBS, 2 mM L-glutamine, 100 units mL^−1^ penicillin, 100 μg mL^−1^ streptomycin, and 20 mM HEPES buffer, all from Gibco Life Technologies) and stimulated with 25 μg *Nme* B16B6 TbpB for 96 hours (at 37°C, 5% CO_2_). Cytokine and chemokine concentrations were determined from supernatants by multiplex array which were performed by Eve Technologies. The data are shown as either a heatmap of median cytokine response from five to six mice per group, normalized to the highest concentration as 1 and the lowest as 0, or in grouped table format in which each animal value is shown by cytokine.

### Genital tract challenge with *N. gonorrhoeae*

4-6 week old female C57BL/6 and BALB/c mice (purchased from Charles River) were allowed to acclimatize for a week, then vaccinated with 25 μg Ngo MS11 TbpB formulations as described above, and then challenged approximately 3-4 weeks after the final dose. Briefly, stage of the estrus cycle was tracked via evaluation of cell morphology of vaginal lavage samples under light microscopy. Upon entering diestrus (Day −2), mice were given β-estradiol (0.5 mg sub-cutaneous, Sigma-Aldrich, cat. E4389) to “lock in” the estrus stage and antibiotics (intra-peritoneal 0.6 mg vancomycin (BioShop Canada, cat. VAN990.5) and 2.4 mg streptomycin (BioShop Canada, cat. STP101.5), plus 0.04 g/100 mL trimethoprim (Sigma-Aldrich, cat. T7883) in the drinking water *ad libitum*) to control overgrowth of commensal flora. For C57BL/6, the injectable antibiotics were administered 2x on Day −1 and then 1x daily until the end of the experiment, and trimethoprim maintained in drinking water for the duration of the study. For BALB/c, the injectable antibiotics were administered 1x on Day −1 and Day 0; and starting Day 2 both streptomycin (0.5 g/100mL) and trimethoprim (0.04 g/100mL) were maintained in drinking water for the duration of the study. Additional doses of β-estradiol were administered on the day of infection (Day 0) and Day +2 to both mouse lines. Baseline vaginal lavages were collected by pipetting 3 times with 30 μl of PBS++ on Day −1. On the day of the infection, mouse passaged Ngo MS11 grown overnight on GC-Kelloggs at 37°C with 5% CO_2_ was suspended in PBS++ and ~5 μl PBS++ containing ~10^7^ CFU was vaginally instilled. Vaginal lavage samples (15 *μ*l PBS++ diluted in 60 *μ*l) were collected daily and dilutions were plated on GC agar supplemented with Kelloggs and VCNT for selection of *N. gonorrhoeae.* Mice were considered no longer colonized when a minimum of three consecutive lavages had no recovery of *N. gonorrhoeae.* Group sizes for each vaccine group and challenge strain can be found in Table S1. Upon completion of the infection study, mice were humanely euthanized by CO_2_ overdose followed by cardiac punction to draw terminal blood. Terminal serum was stored at −20°C until analysis. Gonococcal studies were performed under the animal use protocol 20011775, approved by the Animal Care Committee at the University of Toronto.

### Nasopharyngeal challenge with *N. meningitidis*

Approximately 6 week old male and female transgenic hCEACAM1+/− (mCeacam1−/−) C57Bl/6 and hCEACAM1+/− (mCeacam1+/+) FvB mice were immunized with 25 μg Nme B16B6 TbpB formulations as described above and then challenged ~3 weeks after the final dose. Overnight lawns of mouse-passaged Nme B16B6 grown on GC-Kelloggs at 37°C 5% CO_2_ was suspended in PBS++ and ~10^7^ CFU in 10 *μ*L was inoculated into the nostrils. Mice were humanely euthanized by CO_2_ overdose 3 days post infection., with terminal serum collected by cardiac puncture and stored at −20°C until analysis. Retrograde nasal lavages were collected by injecting 250 *μ*L PBS++ through the trachea and out the nostrils and stored at −20°C until analysis. Nasal turbinates were swabbed and samples grown on GC agar supplemented with Kelloggs and VCNT overnight at 37°C 5% CO_2_ for selection of *N. meningitidis.* Meningococcal studies were performed under the animal use protocol 20011319, approved by the Animal Care Committee at the University of Toronto.

### Protein ELISA

MS11 and B16B6 TbpBs were cloned into a His-Bio-MBP-Tev vector and transformed into *E. coli* strain ER2566. *E. coli* strains were grown overnight in 10 mL cultures of ZYP-5052 autoinduction media containing 100 *μ*g/mL ampicillin at 37°C with shaking. Lysates were obtained by centrifuging the *E. coli* cultures (3,000 g × 5 minutes) and resuspending the pellets in buffer (50 mM NaHP_2_PO_4_, 300 mM NaCl, 10 mM imidazole, pH 8.0). 1 mm glass beads were added to the suspension and cells were disrupted by shaking in a cell disruptor. Homogenate was centrifuged for 20 min at 4°C at 14,000 × g. Supernatant as diluted 1:10 in PBS and added to 384 well ELISA plates (VWR, cat. CA62409-064) that had been previously coated with 1 *μ*g/mL NeutrAvidin (Thermo Fisher, cat. A2666, coated overnight at 4°C) and blocked with 5% bovine serum albumin (BSA, BioShop Canada, cat. ALB001). Protein lysate was coated for 2 hours at room temperature, washed, and then plates were blocked again with 5% BSA for 1 hour. Serum was added overnight at 4°C at dilutions ranging between 1:2,000 and 1:50,000 dilution, and nasal/vaginal samples were added neat. For IgG and IgA ELISAs, standards were utilized by coating wells with goat anti-mouse IgG, IgM and IgA (Southern Biotech #1010-01), followed by addition of serially diluted purified mouse IgG (Abcam, ab37355-1) and IgA (BD Pharmigen, cat. 553478). The following day, plates were washed and relevant secondary antibody was added at 1:5,000 to 1:10,000 dilution. Secondary antibodies used included peroxidase conjugated goat anti-mouse IgG H&L (Jackson ImmunoResearch, cat. 115-035-003), peroxidase conjugated goat anti-mouse IgA (Southern Biotech, cat. 1040-05), alkaline phosphatase conjugated goat anti-mouse IgG1, IgG2a, IgG2b, IgG2c and IgG3 (Jackson ImmunoResearch, cat. 105-055-205, 105-055-206, 105-055-207, 105-055-208, 105-055-209 respectively). Secondary antibody was incubated at room temperature for two hours. For HRP conjugates, plates were developed with KPL SureBlue TMB Microwell Peroxidase Substrate (SeraCare, cat. 5120-0077) and reactions were quenched with 2 N H_2_SO_4_ and read at 450/570 nm. For AP conjugates, plates were developed with BluePhos Substrate (SeraCare, KP 5120-0059) and read at 620 nm.

### Serum bactericidal assays

Mouse-passaged MS11 was grown overnight on GC agar supplemented with Kellogg’s at 37°C with 5% CO_2._ The next day, bacteria was re-streaked onto GC agar supplemented with Kellogg’s and 100 *μ*mol/L deferoxamine mesylate salt (Sigma, cat. D9533) for 4 hours. Bacteria were collected off the plates using Dacron swab, resuspended in RPMI (Wisent, cat. 350-000) and OD_550_measured. Assays were set up in 40 μl total volume using ~500 bacteria suspended in RPMI with 2.5% normal human serum (NHS; Sigma, cat. H4522, lot. SLCD1946) and 2-fold serial dilutions of heat-inactivated terminal mouse serum. The dilution at which 50% killing was observed relative to no antibody control (RPMI with NHS only) in 60 minutes was reported as the SBA titre.

### Statistical Analysis

All statistical analysis was performed using GraphPad Prism 10.3.0 with details included in each corresponding figure legend.

## ACKNOWLEDGEMENTS

We are grateful to Dr. Anthony Schryvers, Dr. Herman Staats and Dr. Cindi Cornelissen for thoughtful conversations during this study, Dr. Nelly Leung for technical support and laboratory management, Dr. Linda Zhong for technical support, and the Division of Comparative Medicine staff at the University of Toronto for animal welfare and colony maintenance.

## REFERENCES

1. Edwards, J.L. and M.A. Apicella, The molecular mechanisms used by Neisseria gonorrhoeae to initiate infection differ between men and women. Clin Microbiol Rev, 2004. 17(4): p. 965–81, table of contents.

2. Eschenbach, D.A. and K.K. Holmes, Acute pelvic inflammatory disease: current concepts of pathogenesis, etiology, and management. Clin Obstet Gynecol, 1975. 18(1): p. 35–56.

3. Hook, E.W. and K.K. Holmes, Gonococcal infections. Ann Intern Med, 1985. 102(2): p. 229–43.

4. Unemo, M. and R.A. Nicholas, Emergence of multidrug-resistant, extensively drug-resistant and untreatable gonorrhea. Future Microbiol, 2012. 7(12): p. 1401–22.

5. St Cyr, S., et al., Update to CDC’s Treatment Guidelines for Gonococcal Infection, 2020. MMWR Morb Mortal Wkly Rep, 2020. 69(50): p. 1911–1916.

6. Rosenstein, N.E., et al., Meningococcal disease. N Engl J Med, 2001. 344(18): p. 1378–88.

7. Wang, N.Y. and A.J. Pollard, The next chapter for group B meningococcal vaccines. Crit Rev Microbiol, 2018. 44(1): p. 95–111.

8. Borrow, R., P. Balmer, and E. Miller, Meningococcal surrogates of protection--serum bactericidal antibody activity. Vaccine, 2005. 23(17-18): p. 2222–7.

9. Borrow, R., et al., The Global Meningococcal Initiative: global epidemiology, the impact of vaccines on meningococcal disease and the importance of herd protection. Expert Rev Vaccines, 2017. 16(4): p. 313–328.

10. Carr, J.P., et al., Impact of meningococcal ACWY conjugate vaccines on pharyngeal carriage in adolescents: evidence for herd protection from the UK MenACWY programme. Clin Microbiol Infect, 2022. 28(12): p. 1649.e1–1649.e8.

11. Maiden, M.C., The impact of protein-conjugate polysaccharide vaccines: an endgame for meningitis? Philos Trans R Soc Lond B Biol Sci, 2013. 368(1623): p. 20120147.

12. Carr, J., et al., ’Be on the TEAM’ Study (Teenagers Against Meningitis): protocol for a controlled clinical trial evaluating the impact of 4CMenB or MenB-fHbp vaccination on the pharyngeal carriage of meningococci in adolescents. BMJ Open, 2020. 10(10): p. e037358.

13. Carr, J.P., et al., Impact of meningococcal ACWY conjugate vaccines on pharyngeal carriage in adolescents: evidence for herd protection from the UK MenACWY programme. Clin Microbiol Infect, 2022. 28(12): p. 1649.e1–1649.e8.

14. Currie, E.G. and S.D. Gray-Owen, Exploring the Ability of Meningococcal Vaccines to Elicit Mucosal Immunity: Insights from Humans and Mice. Pathogens, 2021. 10(7).

15. Read, R.C., et al., Effect of a quadrivalent meningococcal ACWY glycoconjugate or a serogroup B meningococcal vaccine on meningococcal carriage: an observer-blind, phase 3 randomised clinical trial. Lancet, 2014. 384(9960): p. 2123–31.

16. Leong, L.E.X., et al., The genomic epidemiology of Neisseria meningitidis carriage from a randomised controlled trial of 4CMenB vaccination in an asymptomatic adolescent population. Lancet Reg Health West Pac, 2024. 43: p. 100966.

17. McMillan, M., H.S. Marshall, and P. Richmond, 4CMenB vaccine and its role in preventing transmission and inducing herd immunity. Expert Rev Vaccines, 2022. 21(1): p. 103–114.

18. Marjuki, H., et al., Genetic Similarity of Gonococcal Homologs to Meningococcal Outer Membrane Proteins of Serogroup B Vaccine. mBio, 2019. 10(5).

19. Tinsley, C.R. and X. Nassif, Analysis of the genetic differences between Neisseria meningitidis and Neisseria gonorrhoeae: two closely related bacteria expressing two different pathogenicities. Proc Natl Acad Sci U S A, 1996. 93(20): p. 11109–14.

20. Petousis-Harris, H., et al., Effectiveness of a group B outer membrane vesicle meningococcal vaccine against gonorrhoea in New Zealand: a retrospective case-control study. Lancet, 2017. 390(10102): p. 1603–1610.

21. Leduc, I., et al., The serogroup B meningococcal outer membrane vesicle-based vaccine 4CMenB induces cross-species protection against Neisseria gonorrhoeae. PLoS Pathog, 2020. 16(12): p. e1008602.

22. Ladhani, S.N., et al., Use of a meningococcal group B vaccine (4CMenB) in populations at high risk of gonorrhoea in the UK. Lancet Infect Dis, 2024. 24(9): p. e576–e583.

23. Semchenko, E.A., C.J. Day, and K.L. Seib, The Neisseria gonorrhoeae Vaccine Candidate NHBA Elicits Antibodies That Are Bactericidal, Opsonophagocytic and That Reduce Gonococcal Adherence to Epithelial Cells. Vaccines (Basel), 2020. 8(2).

24. Sikora, A.E., et al., A novel gonorrhea vaccine composed of MetQ lipoprotein formulated with CpG shortens experimental murine infection. Vaccine, 2020. 38(51): p. 8175–8184.

25. Price, G.A., M.W. Russell, and C.N. Cornelissen, Intranasal administration of recombinant Neisseria gonorrhoeae transferrin binding proteins A and B conjugated to the cholera toxin B subunit induces systemic and vaginal antibodies in mice. Infect Immun, 2005. 73(7): p. 3945–53.

26. Price, G.A., et al., Gonococcal transferrin binding protein chimeras induce bactericidal and growth inhibitory antibodies in mice. Vaccine, 2007. 25(41): p. 7247–60.

27. Fegan, J.E., et al., Utility of Hybrid Transferrin Binding Protein Antigens for Protection Against Pathogenic Neisseria Species. Front Immunol, 2019. 10: p. 247.

28. Schryvers, A.B., Targeting bacterial transferrin and lactoferrin receptors for vaccines. Trends Microbiol, 2022. 30(9): p. 820–830.

29. Kammerman, M.T., et al., Molecular Insight into TdfH-Mediated Zinc Piracy from Human Calprotectin by Neisseria gonorrhoeae. mBio, 2020. 11(3).

30. Maurakis, S., et al., The novel interaction between Neisseria gonorrhoeae TdfJ and human S100A7 allows gonococci to subvert host zinc restriction. PLoS Pathog, 2019. 15(8): p. e1007937.

31. Jerse, A.E., et al., Estradiol-Treated Female Mice as Surrogate Hosts for Neisseria gonorrhoeae Genital Tract Infections. Front Microbiol, 2011. 2: p. 107.

32. Johswich, K.O., et al., In vivo adaptation and persistence of Neisseria meningitidis within the nasopharyngeal mucosa. PLoS Pathog, 2013. 9(7): p. e1003509.

33. Buckwalter, C.M., et al., Discordant Effects of Licensed Meningococcal Serogroup B Vaccination on Invasive Disease and Nasal Colonization in a Humanized Mouse Model. J Infect Dis, 2017. 215(10): p. 1590–1598.

34. Pajon, R., et al., A native outer membrane vesicle vaccine confers protection against meningococcal colonization in human CEACAM1 transgenic mice. Vaccine, 2015. 33(11): p. 1317–1323.

35. Gulati, S., et al., Immunization against a saccharide epitope accelerates clearance of experimental gonococcal infection. PLoS Pathog, 2013. 9(8): p. e1003559.

36. Liu, Y., et al., Intravaginal Administration of Interleukin 12 during Genital Gonococcal Infection in Mice Induces Immunity to Heterologous Strains of. mSphere, 2018. 3(1).

37. Currie, E.G., Rojas, O., Lee, I., Khaleghi, K., Martin, Al., Gommerman, J. and Gray-Owen, S.D., Protection against Neisseria meningitidis nasopharyngeal colonization relies on antibody opsonization and phagocytosis by neutrophils. [Manuscript submitted for publication]. 2024.

38. Cox, J.C. and A.R. Coulter, Adjuvants--a classification and review of their modes of action. Vaccine, 1997. 15(3): p. 248–56.

39. Awate, S., L.A. Babiuk, and G. Mutwiri, Mechanisms of action of adjuvants. Front Immunol, 2013. 4: p. 114.

40. Korsholm, K.S., et al., T-helper 1 and T-helper 2 adjuvants induce distinct differences in the magnitude, quality and kinetics of the early inflammatory response at the site of injection. Immunology, 2010. 129(1): p. 75–86.

41. Li, H., et al., Cutting edge: inflammasome activation by alum and alum’s adjuvant effect are mediated by NLRP3. J Immunol, 2008. 181(1): p. 17–21.

42. Mata-Haro, V., et al., The vaccine adjuvant monophosphoryl lipid A as a TRIF-biased agonist of TLR4. Science, 2007. 316(5831): p. 1628–32.

43. Casella, C.R. and T.C. Mitchell, Putting endotoxin to work for us: monophosphoryl lipid A as a safe and effective vaccine adjuvant. Cell Mol Life Sci, 2008. 65(20): p. 3231–40.

44. Didierlaurent, A.M., et al., AS04, an aluminum salt- and TLR4 agonist-based adjuvant system, induces a transient localized innate immune response leading to enhanced adaptive immunity. J Immunol, 2009. 183(10): p. 6186–97.

45. Wack, A., et al., Combination adjuvants for the induction of potent, long-lasting antibody and T-cell responses to influenza vaccine in mice. Vaccine, 2008. 26(4): p. 552–61.

46. Higgins, D.A., J.R. Carlson, and G. Van Nest, MF59 adjuvant enhances the immunogenicity of influenza vaccine in both young and old mice. Vaccine, 1996. 14(6): p. 478–84.

47. Knudsen, N.P., et al., Different human vaccine adjuvants promote distinct antigen-independent immunological signatures tailored to different pathogens. Sci Rep, 2016. 6: p. 19570.

48. Klinguer-Hamour, C., et al., DDA adjuvant induces a mixed Th1/Th2 immune response when associated with BBG2Na, a respiratory syncytial virus potential vaccine. Vaccine, 2002. 20(21-22): p. 2743–51.

49. Shibaki, A. and S.I. Katz, Induction of skewed Th1/Th2 T-cell differentiation via subcutaneous immunization with Freund’s adjuvant. Exp Dermatol, 2002. 11(2): p. 126–34.

50. Abara, W.E., et al., Planning for a Gonococcal Vaccine: A Narrative Review of Vaccine Development and Public Health Implications. Sex Transm Dis, 2021. 48(7): p. 453–457.

51. Chow, E.P.F., C.K. Fairley, and F.Y.S. Kong, STI pathogens in the oropharynx: update on screening and treatment. Curr Opin Infect Dis, 2024. 37(1): p. 35–45.

52. Bazan, J.A., D.S. Stephens, and A.N. Turner, Emergence of a novel urogenital-tropic Neisseria meningitidis. Curr Opin Infect Dis, 2021. 34(1): p. 34–39.

53. Cornelissen, C.N., et al., The transferrin receptor expressed by gonococcal strain FA1090 is required for the experimental infection of human male volunteers. Mol Microbiol, 1998. 27(3): p. 611–6.

54. Danve, B., et al., Transferrin-binding proteins isolated from Neisseria meningitidis elicit protective and bactericidal antibodies in laboratory animals. Vaccine, 1993. 11(12): p. 1214–20.

55. Lissolo, L., et al., Evaluation of transferrin-binding protein 2 within the transferrin-binding protein complex as a potential antigen for future meningococcal vaccines. Infect Immun, 1995. 63(3): p. 884–90.

56. Rokbi, B., et al., Evaluation of recombinant transferrin-binding protein B variants from Neisseria meningitidis for their ability to induce cross-reactive and bactericidal antibodies against a genetically diverse collection of serogroup B strains. Infect Immun, 1997. 65(1): p. 55–63.

57. Jamie E. Fegan*, E.A.I., David M. Curran, Dixon Ng, Natalie Au, Elissa G. Currie, Joseph Zeppa, Jessica Lam, Anthony B. Schryvers, Trevor F. Moraes, and Scott. D. Gray-Owen, Bivalent TbpB formulations elicit broad spectrum coverage against Neisseria gonorrhoeae. [Manuscript submitted for publication]. 2024.

58. Schmidt, S.T., et al., The administration route is decisive for the ability of the vaccine adjuvant CAF09 to induce antigen-specific CD8(+) T-cell responses: The immunological consequences of the biodistribution profile. J Control Release, 2016. 239: p. 107–17.

59. Pulendran, B., P. S Arunachalam, and D.T. O’Hagan, Emerging concepts in the science of vaccine adjuvants. Nat Rev Drug Discov, 2021. 20(6): p. 454–475.

60. Pantel, A., et al., A new synthetic TLR4 agonist, GLA, allows dendritic cells targeted with antigen to elicit Th1 T-cell immunity in vivo. Eur J Immunol, 2012. 42(1): p. 101–9.

61. Packiam, M., et al., Mouse strain-dependent differences in susceptibility to Neisseria gonorrhoeae infection and induction of innate immune responses. Infect Immun, 2010. 78(1): p. 433–40.

62. Song, W., et al., Local and humoral immune responses against primary and repeat Neisseria gonorrhoeae genital tract infections of 17beta-estradiol-treated mice. Vaccine, 2008. 26(45): p. 5741–51.

63. Sellers, R.S., et al., Immunological variation between inbred laboratory mouse strains: points to consider in phenotyping genetically immunomodified mice. Vet Pathol, 2012. 49(1): p. 32–43.

64. Studier, F.W., Stable expression clones and auto-induction for protein production in E. coli. Methods Mol Biol, 2014. 1091: p. 17–32.

