## supplemental figures for "Adjuvant-dependent impacts on vaccine-induced humoral responses and protection in preclinical models of nasal and genital colonization by pathogenic Neisseria"

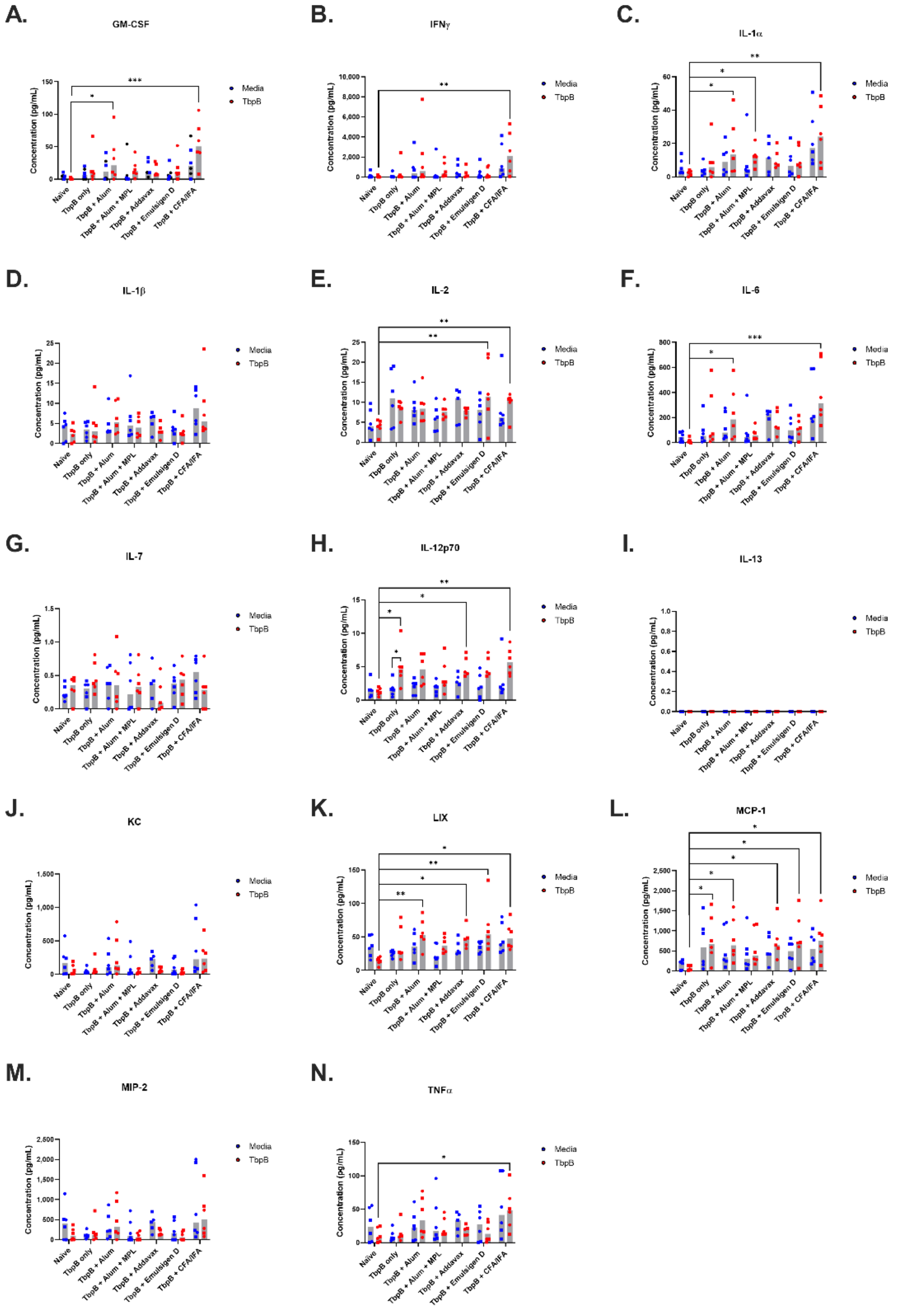


**Figure S1. Cytokine responses from TbpB vaccinated C57BL/6 mouse splenocytes stimulated with TbpB.** Splenocytes from mice vaccinated three times with TbpB plus the indicated adjuvant were stimulated *ex vivo* with purified TbpB protein or media-only for 96 hours. Supernatants analyzed for multiple cytokines and chemokines. Data points represent an individual animal's cytokine concentration to the indicated stimulator (media, blue; TbpB, red). Shaded gray bars represent the median and symbol shape indicates animal sex (square, female; circle, male). Significance was determined using Dunn's non-parametric multiple comparison with control for joint ranks. ***, p<0.001; **, p<0.01; * p<0.05

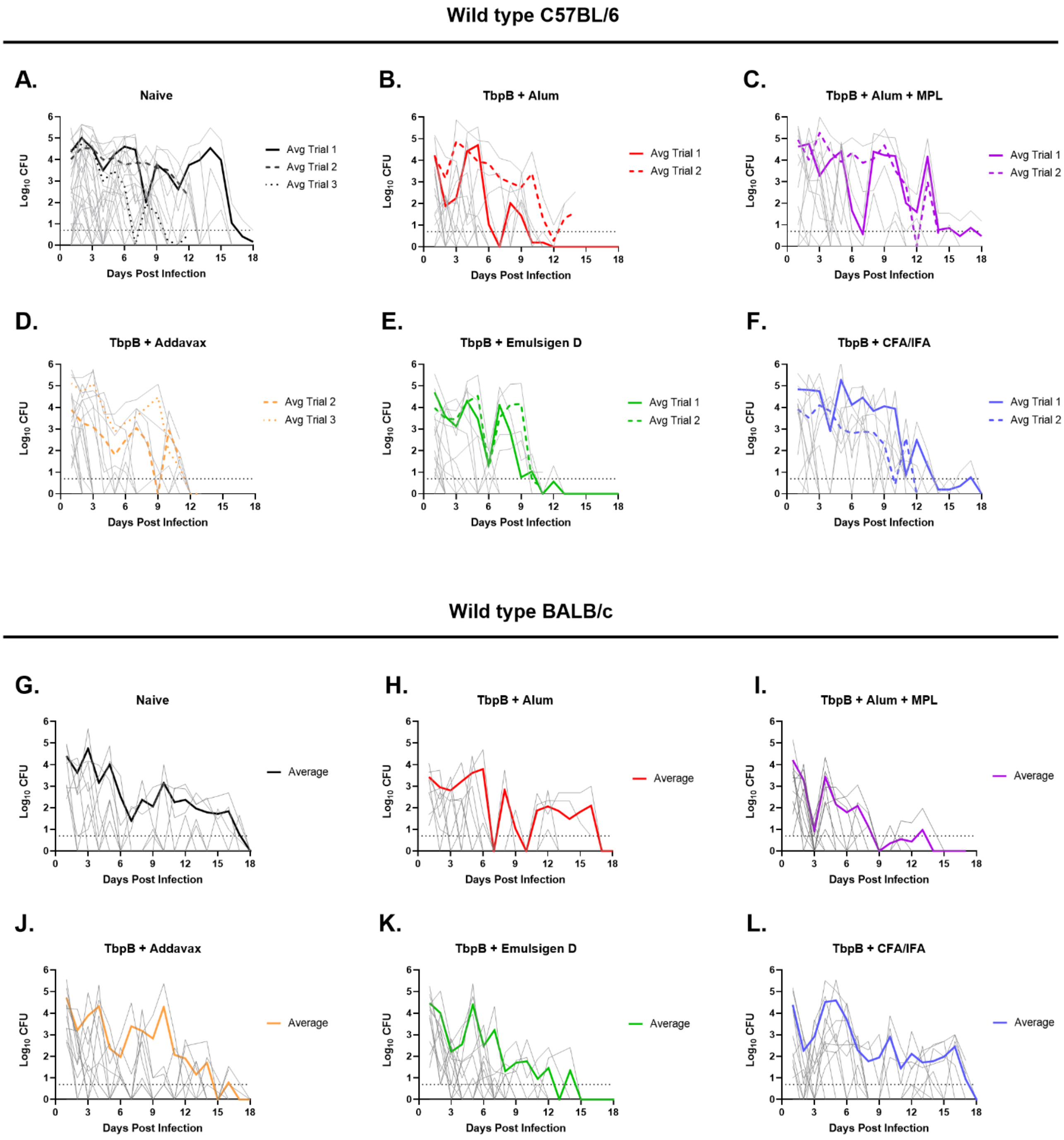


<u>Figure S2</u>: **Gonococcal recovery from the lower genital tract of naïve and TbpB-immunized animals**. Gonococci cultured from daily vaginal lavages of **A-F:** C57BL/6 and **G-L:** BALB/c mice that were vaccinated 3x with the indicated TbpB-formulation is shown. Individual animals plotted in gray; thick lines depict the average CFU from each day for each trial, where applicable. N=6-12 per trial; see Table S1 for detailed breakdown.

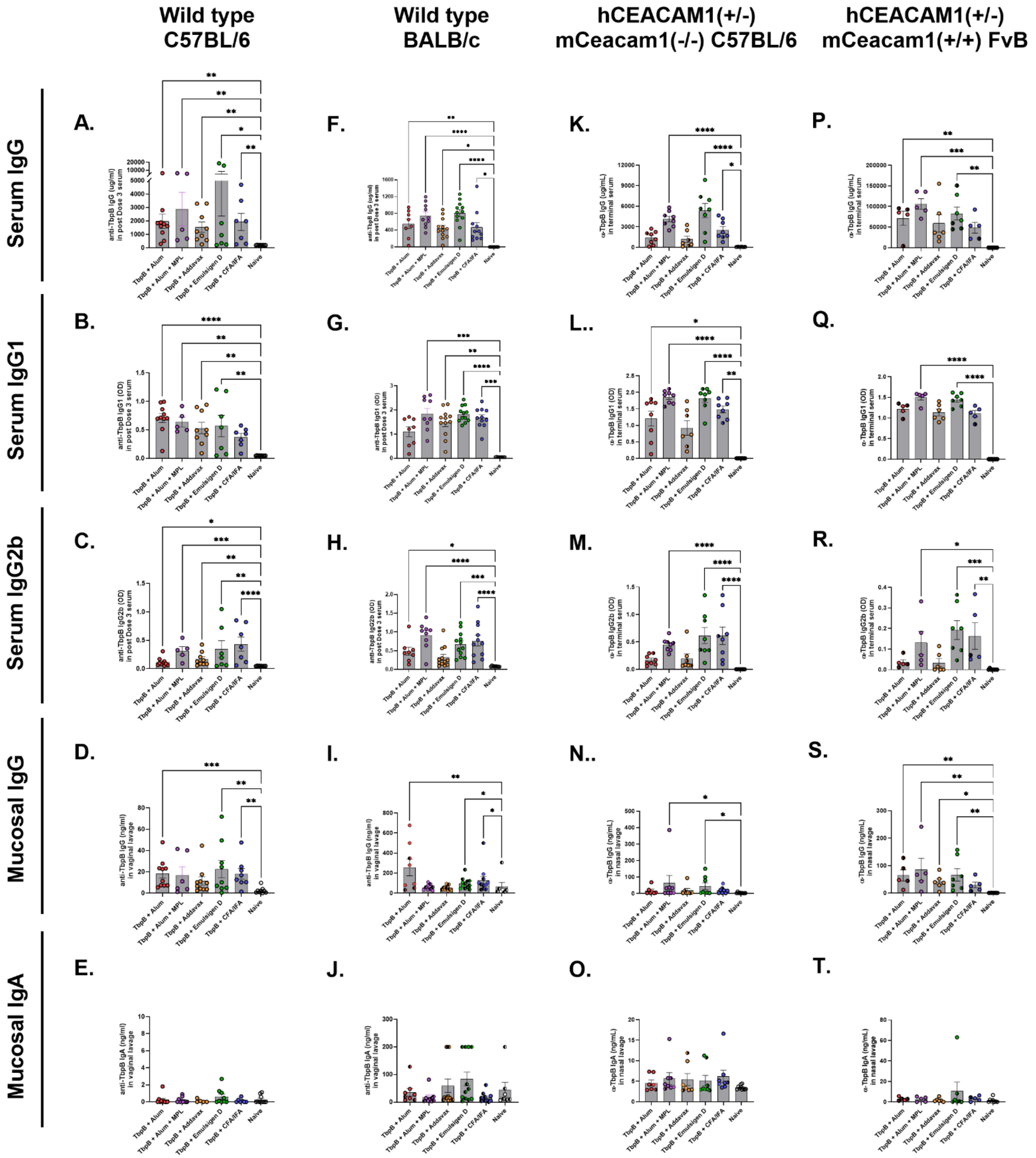


**<u>Figure S3:</u> Immune responses elicited by the different TbpB-based formulations in wild type and transgenic mouse lines used in challenge studies. A, F:** Serum IgG; **B, G:** Serum IgG1; **C, H:** Serum IgG2b; **D, I:** Vaginal IgG; **E, J:** Vaginal IgA in pre-challenge serum and pre-challenge vaginal lavages of wild type C57BL/6 and BALB/c mice immunized 3x with MS11 TbpB with the indicated adjuvant. **K, P:** Serum IgG; **L, Q:** Serum IgG1; **M, R:** Serum IgG2b; **N, S:** Nasal IgG; **O, T:** Nasal IgA in terminal serum and lavages collected from infected hCEACAM1+/- (mCeacam1-/-) C57BL/6 and hCEACAM1+/- (mCeacam1+/+) FvB transgenic mice immunized 3x with B16B6 TbpB with the indicated adjuvant. Antibody levels quantified by ELISA using plates coated with the immunizing antigen. Bars represent mean; error bars depict standard error. N=5-12 animals per group. Trial 2 samples used for quantification for wild type C57BL/6, Trial 1 for other lines; see Table S1 for detailed breakdown of N. Non-parametric Kruskal-Walllis test with Dunn's multiple comparison was performed to compare each TbpB immunized group to the unvaccinated control group. ****, $p<0.0001$; ***, $p<0.001$; **, $p<0.01$; * $p<0.05$. Only significant p-values shown on graphs.
