## supplemental table for "Adjuvant-dependent impacts on vaccine-induced humoral responses and protection in preclinical models of nasal and genital colonization by pathogenic Neisseria"

**Table S1: Number of animals used in gonococcal and meningococcal challenge trials.**

| Mouse Line | Group | Trial 1 | Trial 2 | Trial 3 | Total |
|---|---|---|---|---|---|
| Wild type C57BL/6 | Naïve | 9 | 9 | 12 | N=18 (Trial 1+2)<br>N=19 (Trial 2+3)<br>N= 30 (Trial 1+2+3) |
| | TbpB + Alum | 7 | 10 | | 17 |
| | TbpB + Alum + MPL | 7 | 6 | | 13 |
| | TbpB + Addavax | | 8 | 9 | 17 |
| | TbpB + Emulsigen D | 7 | 9 | | 16 |
| | TbpB + CFA/IFA | 7 | 8 | | 15 |
| Wild type BALB/c | Naïve | 9 | | | 9 |
| | TbpB + Alum | 8 | | | 8 |
| | TbpB + Alum + MPL | 11 | | | 11 |
| | TbpB + Addavax | 12 | | | 12 |
| | TbpB + Emulsigen D | 12 | | | 12 |
| | TbpB + CFA/IFA | 12 | | | 12 |
| hCEACAM1+/- (mCeacam1-/-) C57BL/6 | Naïve | 9 | 6 | | 15 |
| | TbpB + Alum | 8 | 6 | | 14 |
| | TbpB + Alum + MPL | 8 | 6 | | 14 |
| | TbpB + Addavax | 7 | 7 | | 14 |
| | TbpB + Emulsigen D | 8 | 6 | | 14 |
| | TbpB + CFA/IFA | 8 | 6 | | 14 |
| hCEACAM1+/- (mCeacam1+/+) FvB | Naïve | 9 | 9 | | 18 |
| | TbpB + Alum | 5 | 4 | | 9 |
| | TbpB + Alum + MPL | 5 | 6 | | 11 |
| | TbpB + Addavax | 6 | 3 | | 9 |
| | TbpB + Emulsigen D | 5 | 6 | | 11 |
| | TbpB + CFA/IFA | 7 | 6 | | 13 |
